# Global analysis of non-animal peroxidases provides insights into the evolutionary basis of this gene family in green lineage

**DOI:** 10.1101/851881

**Authors:** Duchesse Lacour Mbadinga Mbadinga, Qiang Li, Philippe Ranocha, Yves Martinez, Endymion D. Cooper, Christophe Dunand

**Author notes:** Authors have contributed equally to this work (Duchesse Lacour Mbadinga Mbadinga, Qiang Li). **Corresponding author: Christophe Dunand**.

## Abstract

The non-animal peroxidases belong to a superfamily of oxidoreductases that reduce the hydrogen peroxide and oxidize numerous substrates. Since their initial characterization in 1992, several advances have provided an understanding into the origin and evolutionary history of this family of proteins. Here, we report for the first time an exhaustive evolutionary analysis of non-animal peroxidases using integrated *in silico* and biochemical strategies. Thanks to the availability of numerous genomic sequences from many species belonging to different kingdoms together with expert and exhaustive annotation of peroxidase sequences centralized in a dedicated database, we have deepened our understanding of the evolutionary process underlying non-animal peroxidases through phylogenetic reconstructions. We analysed the distribution of all non-animal peroxidases in more than 200 eukaryotic organisms *in silico*. First, we show that the presence or absence of non-animal peroxidases can be correlated with the presence or absence of certain organelles or with specific biological processes. Examining a wide range of organisms, we confirmed that ascorbate peroxidases (APx) and cytochromes c peroxidases (CcP) were detected respectively in chloroplast and mitochondria containing organisms. Plants, which contain both organelles, are an exception and contain only APxs without CcP. Class III peroxidases (CIII Prx) were only detected in plants and Class II peroxidases (CII Prx) in fungi related to wood decay and plant degradation.

Moreover, we demonstrate that biochemical activities (APx, CcP and CIII Prx) assayed in protein extracts obtained from 30 different eukaryotic organisms strongly support the distribution of the sequences resulting from our *in silico* analysis. The biochemical results confirmed both the presence and classification of non-animal peroxidase encoding sequences.

## Introduction

The increase of oxygen level, 2.3 billion years ago at the beginning of the Proterozoic, forced the ancestral organisms to adapt to oxidative stress (Becker *et al.*, 2004). The reactive oxygen species (ROS) produced are necessary to cells, but toxic, at high levels, and are consequently managed by an ROS gene network (Mittler *et al.*, 2004). Peroxidases are components of this network that regulate ROS homeostasis through oxidation-reduction reactions using hydrogen peroxide (H_2_O_2_) as an electron acceptor along with different oxidized substrates as electron donors. The term “peroxidase” encompasses two major protein families that differ based on in the presence or absence of a haem, namely "haem peroxidase" and "non-haem peroxidase". The haem peroxidases are the most diverse group in terms of taxonomic distribution, and the non-animal peroxidase family belongs to this group (Passardi *et al.*, 2007a). This family was first described by Welinder in 1992 based on structural homology (Welinder, 1992) and comprises three peroxidase classes: Class I (CI Prx), Class II (CII Prx) and Class III (CIII Prx) (Fig 1A). This large superfamily is regrouped under a unique PFAM (PF00141), which described the conserved peroxidase domain. However, this grouping does not discriminate between the members of these three classes and can produce mis-annotation which need to be corrected by manual annotation.

**Fig. 1.**
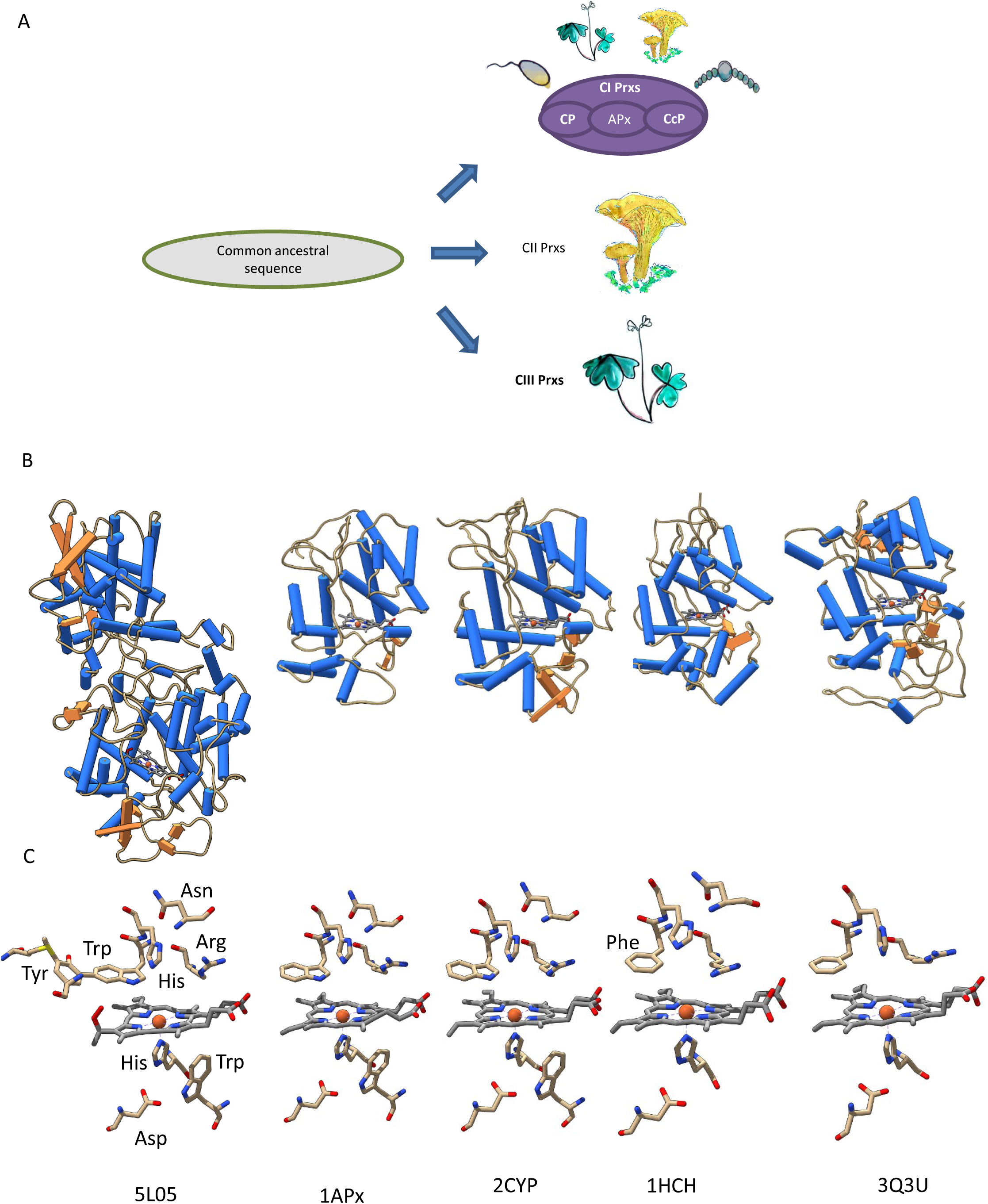
Non-animal peroxidases: 3D structures and catalytic domain are conserved. Class composition and taxonomic distribution (A). The available tertiary structure of 3 CI Prxs (5L05 from *Burkholderia pseudomallei*, 1APx from *Pisum sativum* and 2CYP from *Saccharomyces cerevisiae*), 1 CIII Prx (1HCH from *Armoracia rusticana*) and 1 CII Prx (3Q3U from *Trametes cervina*) were visualized and compared using Swiss-PdbViewer. The 3D models show strong conservation in the number of α-helixes (blue) and their overall spatial organisation (B). Moreover, the key residues necessary for haem binding and electron transfer are also highly conserved amongst among the 3 Prx Classes (C).

The CI Prxs are widely distributed in prokaryotes and eukaryotes and were originally described as intracellular peroxidases (lacking signal peptide). They are composed of an unglycosylated haem, are devoid of calcium ions, and contain no disulphide bridges. This class is the most widespread of the peroxidases and is subdivided into four subclasses: Catalase Peroxidases (CPs), Cytochrome c Peroxidases (CcPs), Ascorbate Peroxidases (APxs) and Hybrid Ascorbate Peroxidase-Cytochrome c Peroxidases (APx-CcPs). CPs are predominantly found in bacteria and only occasionally in other organisms due to horizontal gene transfer (Passardi *et al.*, 2007b). All CPs are characterized by the presence of two independent peroxidase domains. CcPs are present in organisms with mitochondria but are absent from plants. CcPs catalyse the oxidation of ferricytochrome c in the presence of H_2_O_2_ (Finzel *et al.*, 1984) in order to remove toxic oxygen generated during the mitochondrial respiratory process (Kwon *et al.*, 2003). APx are specific to chloroplast-containing organisms with the notable exception of cyanobacteria (Miyake *et al.*, 1991). They catalyse the oxidation of ascorbate in the presence of H_2_O_2_ for photoprotection and regulation of photooxidative stress (Maruta *et al.*, 2010). The last CI Prx subclass corresponds to proteins with common characteristics to both APx and CcP, and has been named hybrid Ascorbate Peroxidase-Cytochrome c Peroxidase (APx-CcP). A predominant example of this class is found in *Leishmana major* for which both APx and CcP activities have been detected (Adak and Datta, 2005).

The CII Prxs are described as secreted fungal lignin peroxidases. They belong to large multigenic families and can oxidize molecules with high redox potential such as lignin. CII Prxs include lignin peroxidases (LiP), manganese peroxidases (MnP), and versatile peroxidases (VPs), so far only described in fungi involved in the degradation of wood (Welinder et al. 1992). More recently, six other classes have been identified in Ascomycetes (Asco class II type A, B and C) and in Basidiomycetes (Basidio class II type A, B and C) (Mathé *et al.*, 2019). These new classes have been primarily found in plant pathogenic and plant degrading fungi.

The CIII Prxs also known as plant secreted peroxidases have been described as potential multifunctional proteins, which can participate in auxin metabolism, cell wall elongation, stiffening and protection against pathogens (Hiraga *et al.*, 2001; Passardi *et al.*, 2004). However, the precise *in vivo* role of any given single peroxidase is still marginal, primarily because of the wide range of peroxidase substrates (lignin subunits, lipid membranes, and some amino acid side chains (Lazzarotto *et al.*, 2011)) and the probable functional redundancy of this large multigenic family that is highly conserved in terms of amino acid sequence. CIII Prxs have been detected in both green algae that predate land colonization and in land plants, but they are absent from Chlorophyte algae (Mathé *et al.*, 2010; Passardi *et al.*, 2007a). This family was subjected to numerous genes, and segmental or genome-wide duplications leading to a large increase in copy numbers from the probable first non-aquatic organisms (Bryophyte). For example, late diverging plants such as *Arabidopsis thaliana* (Tognolli *et al.*, 2002), and *Oryza sativa* (Passardi *et al.*, 2004) contain high copy numbers generated by gene duplications (73 and 138, respectively). These explosive duplications make the genes ideal markers for the mapping of evolutionary lineages in plants.

Unlike CI Prxs, CII and CIII Prxs have more complex structures marked for several glycosylations, four to five disulfide bridges, two calcium binding sites, and one signal peptide (Martinez, 2002). The genes encoding these two classes containing several introns at conserved positions and were subjected to numerous recent duplication events. Gene duplications, together with intronic sequences, play important roles in organism evolution. Intronic sequences are required for various processes such as alternative splicing, translation efficiency and contain regulatory sequences, encoding other genes or non-coding RNAs (Fawcett *et al.*, 2012). The sequences of new paralogs may diverge in regulatory regions, leading to shifts in expression patterns (subfunctionallisation), while divergence in coding regions may result in acquisition of new functions (neofunctionallisation, (Xu *et al.*, 2012). If the duplicated sequences have no additional function, the accumulation of mutations will rapidly lead to pseudogenisation.

To reveal the emergence and analyse the evolution of the CIII Prxs in green plant lineage, we annotated the CI Prxs (APx, CcP and CP) and CIII Prxs families from publically available genomic and expression data. Particular attention was paid to cover most of the green lineage from the early to late divergent plants. In parallel, we confirmed the presence and absence of the three classes of proteins based on APx, CcP and CIII Prxs enzymatic activities. Finally, we focussed on the evolution of the CIII Prx copy number in the green lineage, and in particular, the CIII Prxs from *Spirogyra sp* was analysed.

## Results and Discussion

### The 3D structure is strikingly conserved between CI, CII and CIII Prxs

The three subclasses CPs, APxs, and CcPs which belong to the CI Prxs have very similar tertiary structures, including a conserved number of α-helix domains and overall organization (Figure 1B). The CPs which contain two independent peroxidase domains, consequently as many α-helices. In addition, residues proximal to the heam such as arginine, as well as distal residues such as histidine, are conserved at the same relative positions among the 3 CI Prxs subclasses (Figure 1C). These structural similarities in term of common heam, iron, key residues and structure conservations, strongly suggest that the CPs, APxs, and CcPs originated from a common ancestral sequence. In addition, the comparison between the tertiary structures of the three classes part of the non-animal peroxidases (CI, CII and CIII Prxs) revealed similarly shared common structural elements including heam, iron, and key residues and structural conservations, strongly suggesting that the three Prx classes could have originated from the same ancestral sequence (Figure 1) (Lazzarotto *et al.*, 2015; Passardi *et al.*, 2007a).

Although CII Prxs are not the primary focus of this study, it was necessary to gain insight into the evolution of this class in order to obtain a more global appreciation of the evolution of all non-animal peroxidases. The high duplication rate of CII Prxs, as well as and the absence of CII Prxs in organisms like Chytrids at the base of fungi and even in some species of basidiomycetes, hinder the clear determination of the evolutionay basis behind this class of peroxidases (Mathé *et al.*, 2019). However, the functional similarity and tertiary structure conservation of CII Prxs with CI Prxs suggest that the CII Prx sequences could have derived from CI Prxs. Therefore, it is likely that the ancestor of the CII Prxs had duplicated and emerged independently during the separation of Basidiomycetes and Ascomycetes to give rise to these different subclasses of CII Prxs (Mathé *et al.*, 2019).

### CI, CII and CIII Prxs have specific taxonomic distribution

Several taxonomic distributions of CI, CII and CIII Prxs have already been described (Lazzarotto *et al.*, 2015; Passardi *et al.*, 2007a), but more recent advances in available comprehensive genomic resources (Mukherjee *et al.*, 2018), allowed us to perform exhaustive data-mining and expert sequence annotation in order to define the evolutionary basis giving rise to the actual taxonomic distribution of the non-animal Prxs in the different kingdoms.

To do this, we performed data mining for peroxidase sequences and organelle gain or loss events from different eukaryotic taxonomic groups that yielded a fine distribution of the different classes (Figure 2, Table 1 and Sup Table 1). We employed two complementary strategies to perform exhaustive and expert data mining. First, NCBI, JGI and 1KP databases were queried using the PFAM annotation (PF00141). As this PFAM encompasses the entire set of haem containing peroxidases (animal and non-animal peroxidases), it allowed us to collect most of the non-animal peroxidase sequences, but did not allow to discriminate between the different classes and subclasses. To complete this approach, we used a homology-based strategy to determine the class/subclass memberships and to detect new sequences not already annotated. All the sequences detected were available from the RedoxiBase (Savelli *et al.*, 2019).

**Table 1.**
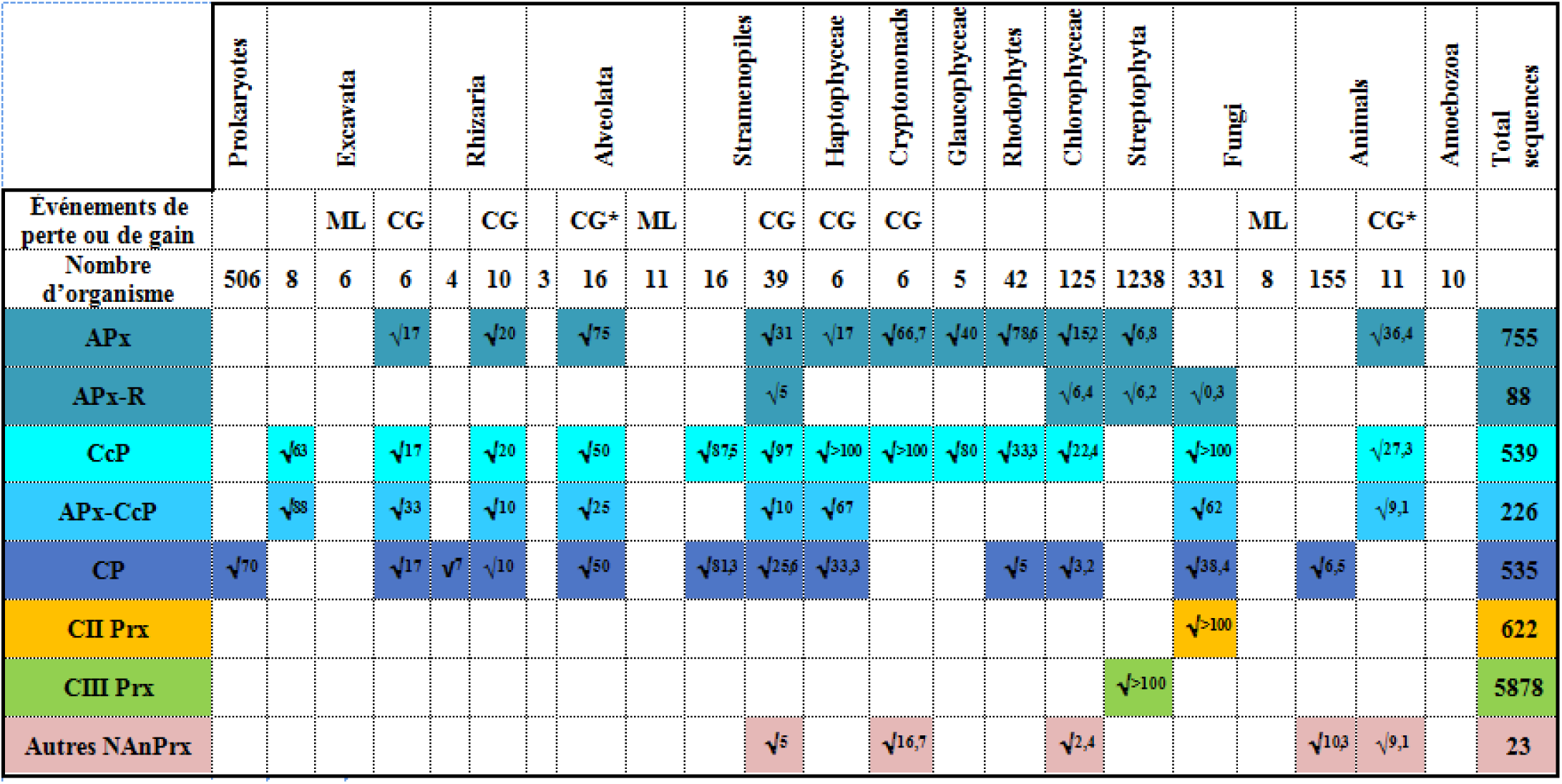
Taxonomic distribution of non-animal haem peroxidases (CI, CII, CIII Prxs) in prokaryotic and eukaryotic organisms. The blue boxes together represent the class I peroxidases (CI Prxs: APx, APr-R, CcP, APx-CcP and CP) in yellow, Class II peroxidases (CII Prxs), and in green, those of class III (CIII Prxs). The presence of this symbol * in certain cells indicates that there is no chloroplast organelle. The chloroplast sequences come from a gene transfer. The superscript numbers represent the % of sequences found in each group and calculated using referenced sequences in Redoxibase.

**Fig. 2.**
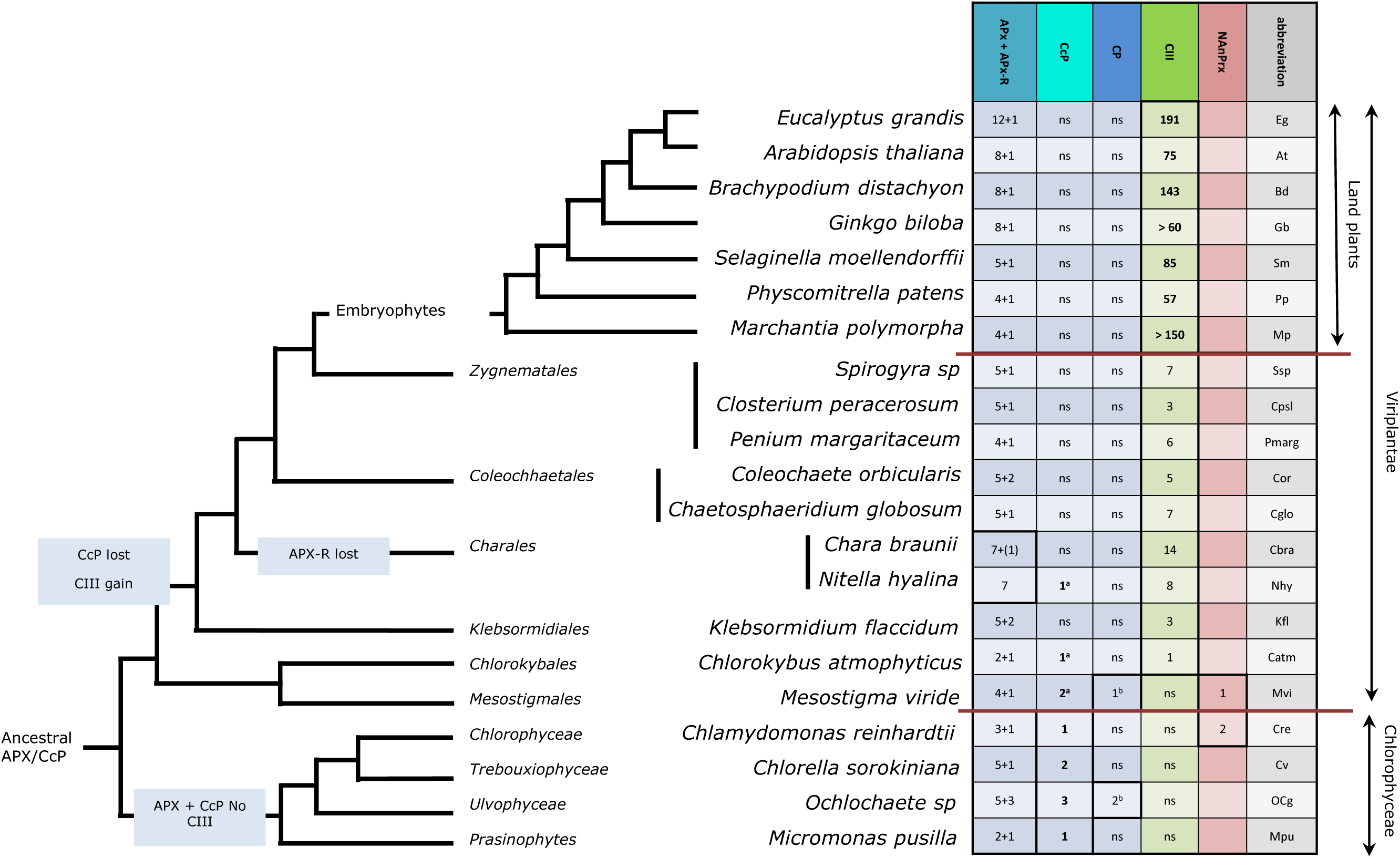
Evolution of CI and CIII Prxs isoform numbers in green lineage. The number of genes encoding CI Prxs (CP, APx and CcP) and CIII Prx was analysed for 21 species from viridiplantae and Chlorophyceae. All the sequences detected and included in the table are available from the RedoxiBase (www.peroxibase.toulouse;inra.fr). Other: other non-animal Prx, ns: no sequence found. Marginal unexpected detection of CcP and CP sequences due to putative sample contamination (^a^, hits only with chlorophyceae), and Lateral Gene Transfer (^b^, LGT)

Thanks to the exhaustive collection and analysis of sequences, we have obtained a fine distribution of non-animal peroxidases. (Figure 2 and Table 1). Our results confirm and strongly support that the concept that APx sequences are only detected in the eukaryotic groups that having acquired and maintained chloroplasts. In the other way around, all the chloroplastic containing organisms analysed (based on available genome sequences) possessed between one to ten APx sequence in the Angiosperms (Table 1). A more divergent sequence, APx-related (APx-R) proteins have only been detected in the green lineage (Chlorophyceae and Streptophyta). Only two diatoms (*Phaeodactylum tricornutum* and *Thalassiosira pseudonana*) and one early divergent fungus (*Gonapodya prolifera*) possessed APx-R encoding sequences, which may have resulted from lateral gene transfers from a green lineage organism (Table 1). Furthermore, the APx-R could have emerged more recently in the green lineage from an existing APX (neofunctionalisation). In addition, CcPs are present in all eukaryotic organisms but absent from Streptophytes and organisms that have undergone a mitochondrial loss event such as Microsporida (obligate intracellular parasite fungi) or which have developed parasitic style of life (*Apicomplexa* animal parasites, *Pyromyces* commensal organisms of the herbivore gut) (Figure 2 and Table 1). Interestingly, we can notice that all the organisms having acquired a chloroplast, have preserved their mitochondria. On the other hand, a loss of mitochondria is never associated with a gain of chloroplast. The majority of the detected CP sequences belong to the prokaryotes (bacteria) while the rest was distributed punctually in several eukaryotic genomes due to probable lateral gene transfer (Passardi *et al.*, 2007a)(Table 1). We found that all CI Prxs are distributed throughout the eukaryotic kingdom confirming the hypothesis of a common ancestral sequence for the different members of the CI Prx subclasses. The APx-CcP hybrid sequences were detected in several kingdoms without taxonomic relations (Table 1). The APx-CcP have conserved key residues from the Class I but are clustered together within the kingdoms meaning that they diverged independently from an ancestral APx or CcP sequence.

Next, with the use of a larger dataset, we confirmed and supported, the hypothesis that APxs are specific to chloroplast containing organisms, and that CcPs are specific to mitochondrial containing organisms with the exception of all embryophytes. In the Excavata, two phyla perfectly illustrated these gain or lost events. On the one hand, *Giardia intestinalis*, *Spironucleus barkhanus* and *Spironucleus salmonicida* belonging to the Diplomonadida phylum have lost their mitochondria and lack the CcP sequence. On the other hand, *Euglena mutabilis*, *E. gracilis* and *Eutreptiella gymnastica* part of Euglenida phylum, have gained chloroplasts and contained both APx and CcP sequences.

A marginal presence of APx sequences was found in non chloroplast-containing organisms, likely due to lateral gene transfer (LGT) (Table 1). This phenomenon is most often observed in living water organisms (proximity with algae containing APx) and for symbiotic organisms. *Hydra viridis*, belonging to animals contained a non-animal Prx with both characteristic of APx and traits of CIII Prx. Based on the homology with APx sequences found in green algae, a LGT from Chlorella can be postulated, and that some subsequent evolutionary convergence or divergence occurred, leading to the homology with CIII Prx. We also note the marginal presence of CI Prxs in organisms belonging to animal kingdoms, even though these proteins are known to be generally absent from the animal kingdom. For example, the presence of APx and CcP encoding sequences in the order of Choanoflagellates demonstrates this.

In addition, we observed a lack of CcP for several parasitic organisms containing mitochondria. Among them, is Apicomplexa, a unicellular animal parasite containing a mitochondria and an organelle referred to as the apicoplast (a relic of chloroplasts obtained by secondary endosymbiosis of red algae, (Lim and McFadden, 2010). More than 10 Apicomplexa genomes are available from NCBI and JGI but no Class I has been detected in these organisms. A chloroplast-containing Apicomplexa, *Chromera* (small EST library) was analysed and an APx like protein was found but no CcP.

Regarding the CII and CIII Prxs, the taxonomic distribution is more precise and has recently been described (Mathé *et al.*, 2019). However, we found that it was necessary to include CII and CIII Prxs in our analysis of non-animal peroxidase evolution, because it allowed us to draw a more integrative evolutionary scenario.

Briefly, the set of CII Prxs sequences was only detected in Ascomycetes and Basidiomycetes that interact with living or dead plants. As mitochondrial containing organisms, Ascomycetes and Basidiomycetes contained CcP in two copies in most cases. This is likely not due to recent gene duplication, as the two copies are phylogenetically distant. We observed that most of the fungi containing CII Prxs lack the second CcP copy, which suggest that CII Prx could have originated from one CcP copy. Microsporidia, obligate intracellular parasite fungi, and Pyromyces (Neocallimastigomycota order), a commensal organism of the herbivore gut, have both lost mitochondria during evolution and did not contain CcP sequence (absence confirmed from more than 10 available genomes).

Similarly to the CII Prxs evolutive scenario, CIII Prxs sequences were only found in Streptophytes and are absent from Chlorophyceae. The number of CIII Prx genes has largely increased along the green lineage (from 0 in Chlorophyceae to 191 *E. grandis*, Figure 2 and Table 1). The late diverging plants contain more CIII Prxs copies than early diverging plants due to explosive and recent duplications (tandem (TDs), segmental (SDs) and whole genome duplications (WGDs), (Li *et al.*, 2015), which make the size of CIII family quite variable (Figure 2). Since Streptophytes are the only mitochondria containing organisms lacking CcP, and because CI and CIII Prxs present high level of homology and several conserved key residues, we raise a hypothesis that the ancestral CIII Prx was derived from the ancestral CcP sequence.

Other non-animal peroxidases, not belonging to any of the well described CI, CII and CIII Prxs, are non-specifically distributed. Therefore, they can be considered as the result of an evolutive divergence or convergence from an ancestral CI Prxs.

### In land plants, the CIII Prx activities are greater than those of other plants

APxs, CcP and CIII Prx specific activities have been tested in different species including early to late divergent plants, several fungi, and chloroplast-containing protists (Figure 3).

**Fig. 3.**
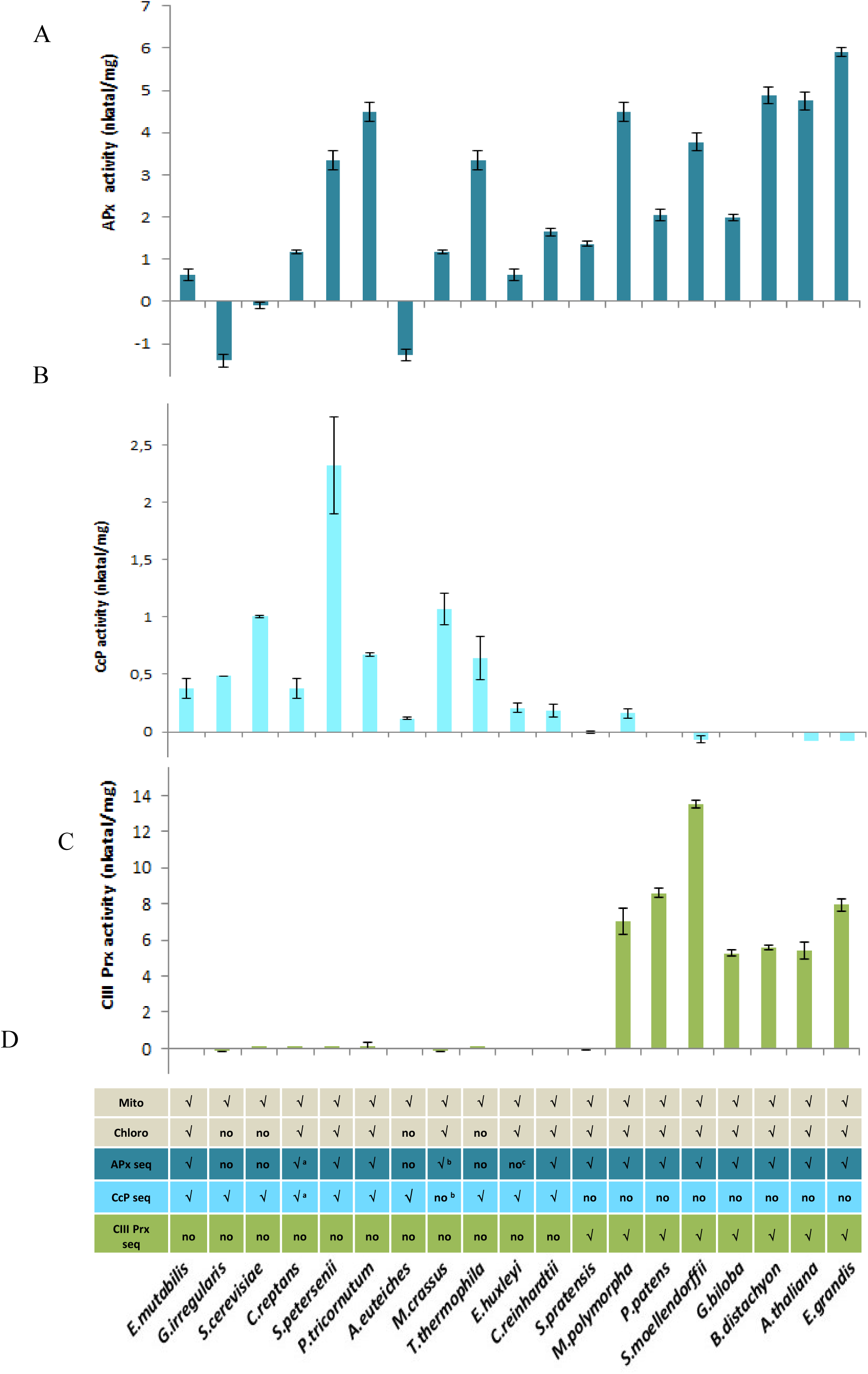
APx, CcP and CIII Prx activities were correlated with the presence of the corresponding encoding proteins. Total protein was extracted from 19 species belonging to green plants, fungi and protists. (A) APx activity was determined by the oxidation of ascorbic acid based on the decrease of the absorbance at 290 nm. (B) CcP activity was measured by monitoring the oxidation of Cytochrome c previously reduced at 550 nm. (C) CIII Prx activity was assayed at 470 nm to the follow the oxidation of guaiacol. (D) Table summarizing, the presence of mitochondria (Mito), CcP encoding sequences (CcP seq), chloroplast (Chloro), APx encoding sequences (APx seq) and CIII Prx encoding sequences (CIII Prx seq)) for the 19 studied species. √: organite, sequences have been detected, no: no organite, or sequence has been detected, ?: no sufficient data to conclude for the presence or the absence of sequences. The enzymatic activities are expressed in nanokatal per mg of total proteins. Activities are the mean of 3 independent assays ± SD.

Significant variations of APx activities were observed between species likely due to the organ variability or different substrate affinity. However, the different activities are in the same order of magnitude between the different organisms from the green lineage tested (Figure 3A), which is in agreement with the conserved number of APx sequences detected with the *in silico* analysis (Figure 2). As expected, no APx activity was detected in non chloroplast-containing organisms. CcP activity was detected in all mitochondria-containing organisms excepted in the eight organisms belonging to the Streptophytes (Figure 2B). This was in agreement with the presence or absence of CcP encoding sequences (Figure 2).

CIII Prxs activity was only detected in the Streptophytes (Figure 2C) which is consistent with our *in silico* analysis. The differences between early divergent plants (*K. flaccidum*, *S. pratensis* and *C. orbicularis*) and late divergent plants were significant (P = 0.01), and the range of values was very high (a ratio of 1000:1 between *C. orbicularis* and *E. grandis*). Between the Angiosperm tested, we observed that the activities were also significantly different possibly due to certain variability in plant ages or stages development. This is in agreement with the finding that the activities of different-aged *A. thaliana* were significantly different (data not shown (Cosio and Dunand, 2010)). Surprisingly, few activity was detected from *K. flaccidum*, *S. pratensis* and *C. orbicularis* protein extracts despite the presence of CIII Prx sequences in these organisms (Figure 2C). This low activity could be due to the poor cell wall protein extractability and/or to the lower affinity for *in vitro* substrate, likely due to the reduced sequence similarity when compared with *A. thaliana* sequences (36 to 42% of identity for the sequences of *K. flaccidum*, *S. pratensis* and *C. orbicularis* when compared with the closest *A. thaliana* orthologous sequence).

In the green lineage, it appeared that the activities of CIII Prxs increased along the green plant evolution while no significant difference of APx between plants or CcP activity has been detected (Figure 3). The explosion of CIII Prx activities can be correlated with the explosion of the gene copy number. Although no direct evidence has been demonstrated yet, this explosion could be related to the variation of environmental oxygen concentration, to the organs and cell wall complexification, to the climatic changes, to the colonization of new biotopes, to the constant appearance of new pathogens and, most recently, to human impact on cultivated plants (Passardi *et al.*, 2004).

### The features of CIII Prx in early divergent plants

In order to confirm the increase in gene number from early to late divergent plants, *Spirogyra* sp. genome and *S. pratensis* RNA-seq data were analysed (Delaux *et al.*, 2015; Van de Poel *et al.*, 2016). Spirogyra belong to the Zygnemophyceae, a close relative of land plants. At least seven independent CIII Prx and five independent APx encoding sequences were detected from *Spirogyra* sp and *S. pratensis*. In parallel, a search of *K. flaccidum* and *Closterium peracerosum* available sequences allowed for the identification of 3 CIII Prx from the two species and seven and six APx, respectively. The CIII Prxs from the two *Spirogyra* sp encode for proteins with regular predicted 3D structure containing 12 alpha-helixes (Sup Figure S1). Although the sequences were not shorter, no predicted signal peptide or C-terminal extension can be detected with various prediction programs (SignalP 4.1 (Petersen *et al.*, 2011), PrediSi (Menne *et al.*, 2000), Signal-3L (Shen and Chou, 2007) and SOSUIsignal (Hirokawa *et al.*, 1998)) for two CIII Prxs of the seven (considering that two sequences are partial). The canonical CIII Prx sequences have eight conserved cysteine (Cys) residues (C1-C8) that are necessary for disulfide bond formation (Figure 2). CIII Prx genes lacking some Cys residues can be found all along the green lineage with a frequency lower than 5 % (41 CIII Prx with missing cysteine for 959 CIII Prx analysed). No genes lacking more than two disulfide bonds have been identified in this study. Despite the absence of data concerning the influence of the lack of disulfide bonds on the protein stability or activity, we could assume that the thermo stability should be decreased due to missing disulfide bonds. In the seven CIII Prxs from *Spirogyra sp*, the cysteines C2 and C3 surrounding the distal histidine are missing while they are present in the CIII Prx from *K. flaccidum* and *C. peracerosum*. The transient expression in tobacco leaves of the native SsPrx03 and SsPrx03 with the two missing cysteines have been performed in *N. benthamiana*. No fluorescence was detected in any cell compartments of *N. benthamiana* leaves transformed with SsPrx03 fused to the TagRFP. However, the cell wall localisation was detected when leaves were transformed with sequence containing the two cysteines C2 and C3 (SsPrx03m1, Figure 4A). However, similar signal was detected in the leaves transformed with CIII Prx from *A. thaliana* (native AtPrx36 and AtPrx36 mutated for the cysteines C2 and C3 (AtPrx36m3)).

**Fig. 4.**
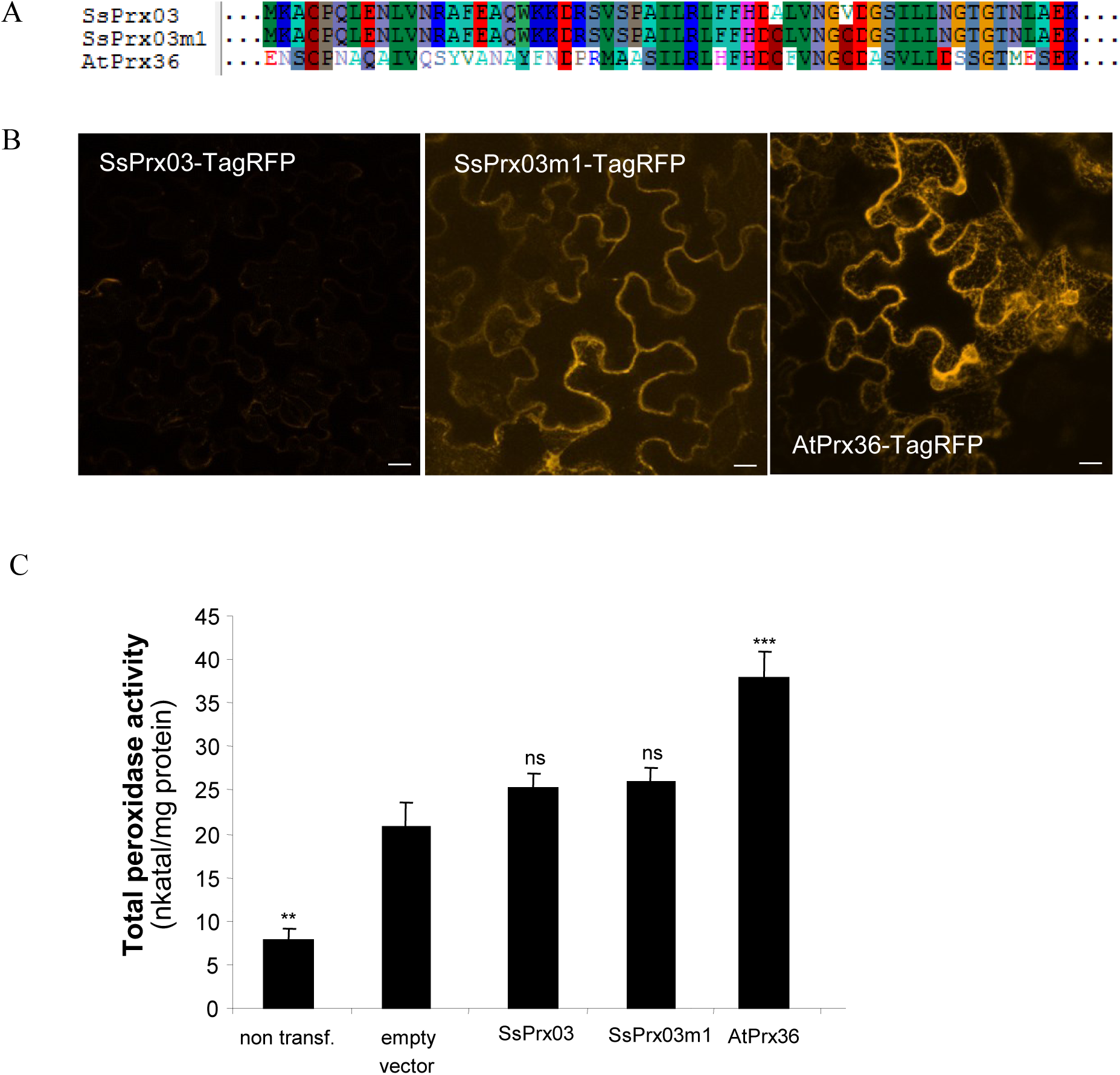
The lack of cysteine C2 and C3 could affect the cell wal localssation of Spirogyra CIII Prx. (A) Alignment of SsPrx03, SsPrx03m1 and AtPrx36 partial region surrounding the conserved distal histidine. (B) Pieces of *Nicotiana benthamiana* leaves transformed with constructs expressing SsPrx03-TagRFP, SsPrx03m1-TagRFP and AtPrx36-TagRFP were cut 48 h post-agroinfiltration. The fluorescence signal was observed with a Leica TCS SP2 Confocal Laser Scanning Microscope (B) Proteins were extracted from *N. benthamiana* leaves transformed with constructs expressing SsPrx03-TagRFP, SsPrx03m1-TagRFP and AtPrx36-TagRFP and assayed for peroxidase activity using guaiacol/hydrogen peroxide as substrate. Enzyme activity values (expressed as nkatal/mg protein) are the mean of at least three replicates ± SD. P-value of one-way anova followed by Tukey’s significance test, (***) P < 0.0001, (**) P < 0.001, (ns) non significanly different to leaves transformed with empty vector.

In parallel, total CIII Prx activity was measured in the infected areas of the leaves. First, the lack of the cysteines C2 and C3 did not significantly affect the CIII Prx activity detected in the leaves of the AtPrx36m3 vs AtPrx36. The addition of the cysteins C2 and C3 in the *Spirogyra sp* sequence did not increase the initial activity (SsPrx03 vs SsPrx03m1) and was not significantly different from the activity detected with the empty vector in both cases (Figure 4B). The low activity detected with SsPrx03, lacking the cysteines C2 and C3, could support the very low activity detected from *Spirogyra sp* fresh material (Figure 3).

### Conclusions

Based on previous studies (Lazzarotto *et al.*, 2015; Passardi *et al.*, 2007a), we have performed exhaustive data-mining and expert sequence annotation. We have greatly increased the number of sequences belonging to underrepresented kingdoms. We conclude that the three CI Prx subclasses (APx, CcP and CP) have originated from the same ancestral sequence initially present in the Neomura (Figure 5). The presence of APx in various photosynthetic organisms (uni, multi and pluricellular organisms) suggests that sequence gains followed independent secondary endosymbiosis. The chaotic distribution of the bifunctionnal CP due to LGT and primarily found in prokaryotic organisms strongly supports the existence of a mono functional ancestral CI Prx. The evolutive scenario leading the modern CII and CIII Prxs appeared to be similar. They probably both emerged independently from an ancient CcP independently in the ascomycetes and bascidiomycetes for the CII Prx and prior to the plant terrestrialisation for the CIII Prxs. The emergence of the CII and CIII Prxs was accompanied by the acquisition of a secretion signal and several cysteine residues allowing disulfide bridge formation, which was coherent with the extracellular activity of these proteins. These common innovations strongly support the common evolutionary basis of CII and CIII Prxs.

**Fig. 5.**
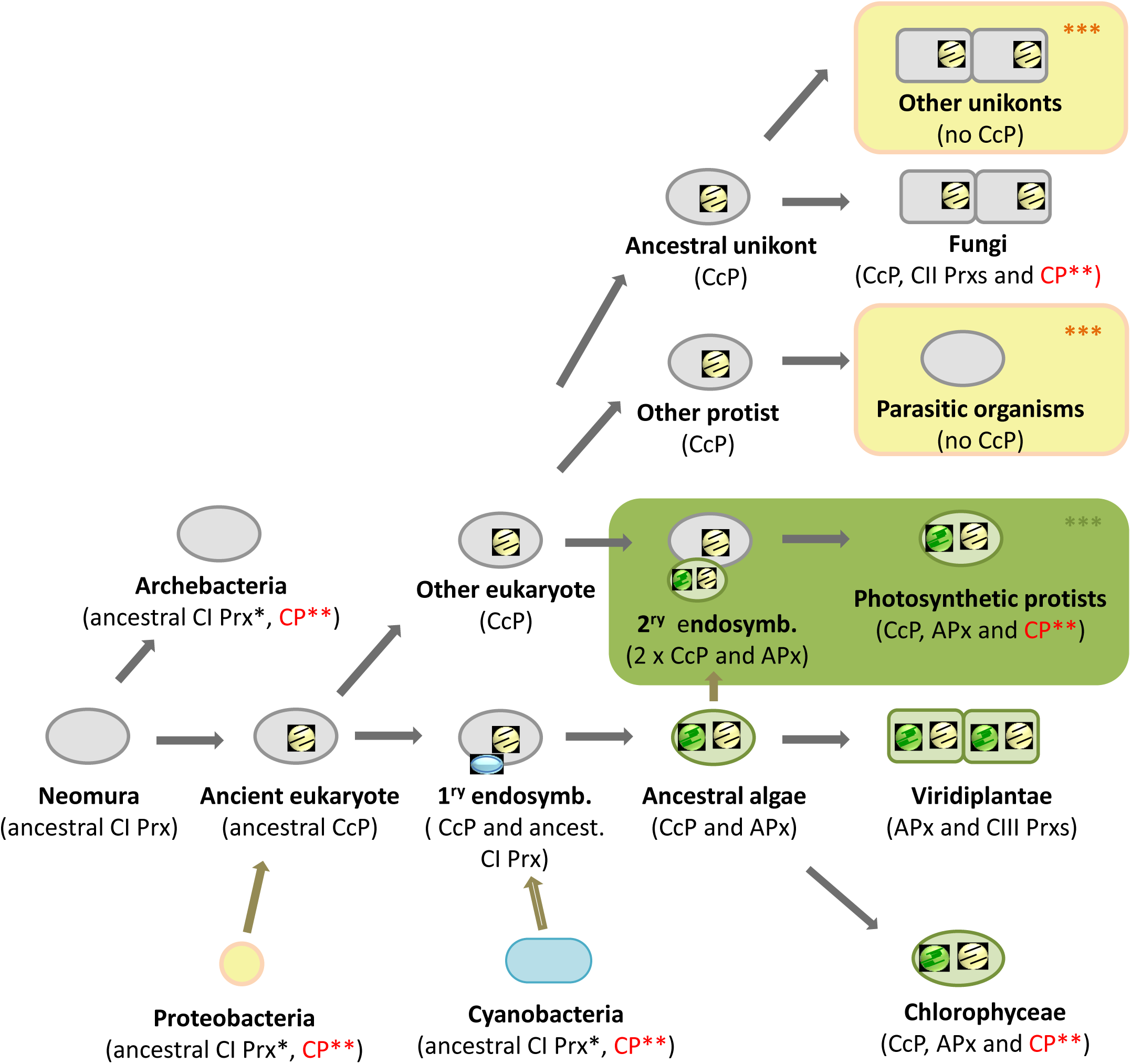
An evolutionary model to explain the current distribution of the different non-animal peroxidase classes. *: sequence lost in the current organisms; **: chaotic distribution resulting from LGT; ***: several independent events.

## Materials and Methods

### Cultivated and collected organisms

The protist (*Euglena mutabilis* (Euglenida), *Chlorarachnion reptans* (Cercozoa), *Synura petersenii (Other Stramenopiles)*, *Moneuplotes crassus* (Alveolates), *Tetrahymena thermophila* (Ciliophora, ciliates), *Emiliania huxleyi* (Isochrysidales)) and the algae (*Chlamydomonas reinhardtii* (*Chlorophyceae*), *Klebsormidium flaccidum*, *Coleochaete. Orbicularis, Spirogyra pratensis*) were provided by the Culture Collection of Algae and Protozoa (CCAP, https://www.ccap.ac.uk/) and cultivated *in vitro* with culture media compositions detailed in sup Table 2. Illustrations of cultivated cultures are visible in sup Fig S2 *Saccharomyces cerevisiae* (*Saccharomycetes*) is baker yeast and protein extraction was made directly without in vitro culture. The two diatoms (*Moneuplotes crassus* and *Phaeodactylum tricornutum* (Bacillariophyta) were provided by the LISBP laboratory and their growth conditions are also describe in sup Table 2.

The Oomycete (*Aphanomyces euteiches*) and the Glomeromycete (*Glomus irregularis* were available from our laboratory and have been cultivated as indicated in sup table 2.The land plants *Marchantia polymorpha*, *Physcomitrella patens*, *Selaginella moellendorffii*, *Gingko biloba*, *Brachypodium distachyon*, *Arabidopsis thaliana*, *Eucalyptus grandis* were available directly from the growth room facility.

### Total protein extraction and enzyme assays of APx, CcP and CIII Prxs

For multicellular organisms, approximately 100 mg fresh material was ground into fine powder with a mortar and pestle under liquid nitrogen. For cell culture, to extract proteins 200 µl extraction buffer containing potassium phosphate buffer (pH 7.0, 50 mM/L), EDTA (5 mM/L), PVPP (16 g/L) was used for extraction. The homogenate was centrifuged at 10, 000 g for 10 min at 4 °C and the supernatant was used for enzyme assays and total protein assay.

The APx activity was determined based on the oxidation of ascorbic acid (ASA). Oxidation was determined based on measurements of the decrease in absorbance at 290 nm between 30 sec to 5 min. one ml reaction mixture contained potassium phosphate buffer (pH 7.0, 50 mM/L), ASA (0.5 mM/L), H_2_O_2_ (0.1mM/L) and 20 µl of protein extract already containing ASA, 2 mM/L. The reaction was initiated with the addition of the of protein extract. A correction was made for the low, non-enzymatic oxidation of ascorbate by H_2_O_2_ (Nakano *et al.*, 1981). The APx unit is defined as the decrease in 1 min of 1 µg total proteins under the above assay conditions.

The CcP activity was measured by monitoring at 550 nm the oxidation of Cytochrome c (from the bovine heart, Sigma-Aldrich) previously reduced. The reduction was carried out by mixing a solution of 0.22 mM Cytochrome c with a 0.5 mM final concentration DTT solution for 15 min at room temperature. After this time, the color of the solution should change from red to pale pink-red for a good reduction. Nevertheless, the reduction must be estimated by spectrometry by measuring the ratio of OD 555 nm / OD 565 nm of an aliquot of the solution diluted 20-fold in the assay buffer. For a good reduction, the ratio should be between 10 and 20. The assay conditions for the CcP activity are a few microliters of protein, 50 mM sodium phosphate buffer pH = 7, 200 μM H_2_O_2_ and 20 μM Cytochrome c in a plastic tub. After adding the proteins or H_2_O_2_, the OD is measured between 10 and 30 seconds.

The CIII Prx activity was assayed in a 1 ml reaction mixture consisting of potassium phosphate buffer (50 mM/L, pH 6.0), 0.125% guaiacol (v/v) and 20 µL of protein extract. The reaction was initiated by adding 125 µL H_2_O_2_ (11 mM/L), and the oxidation of guaiacol was determined based on the increase in A470. One CIII Prx unit is defined as the increase between 1 min and 2 min of 1 µg total proteins under the above assay conditions (Jia *et al.*, 2013).

The concentration of total proteins, used to determine the enzyme activity was detected using Coomassie Protein Assay (Bradford).

### Identification and annotation of CI Prxs and CIII Prx genes in green lineage

The presence or numbers of CI Prxs and CIII Prxs were analysed from various organisms belonging to the green lineage including early and late divergent plants. Expert and exhaustive data-mining was performed from annotated (NCBI, Phytozome v9.1 (http://www.phytozome.net/), NCBI (Benson *et al.*, 1990) and RedoxiBase (Fawal *et al.*, 2013) and non-annotated repositories (1 kP and EST libraries from NCBI). All the annotated peroxidase sequences were all available from the RedoxiBase.

### Subcellular localization of CIII Prx protein SsPrx03

The full-length ORF frame of SsPrx01 (without stop codon) was amplified by PCR from synthetic sequence using 5’-CCGGAATTCATGTCTCTTCTTCCCC-3’ and 5’-CCGCCCGGGAAGATACAAGCAATAC-3’. The PCR product was digested with *EcoR*I and *Sma*I, and cloned into TagRFP-AS-N vector (Evrogen). The SsPrx03 and SsPrx03m1-TagRFP C-terminal fusion was then subcloned (LR reaction, Gateway Technology, Invitrogen) into pEAQ-HT-DEST1 vector (Sainsbury *et al.*, 2009). This construct was confirmed by restriction analysis and sequencing, and transferred into *Agrobacterium tumefaciens* (strain GV3101). The YFP fusion protein pm-yb CD3-1006 (Nelson *et al.*, 2007) was used as a plasma membrane control. Both constructs were co-infiltrated into *N. benthamiana* 30-day old leaves at an optical density of 0.5 at 600 nm. 36 hours after infiltration leaves were detached and used for image acquisition on a Leica TCS SP2 Confocal Laser Scanning Microscope with the following excitation wavelengths: 561 nm for TagRFP and 514 nm for YFP.

### SsPrx03-TagRFP, SsPrx03m-TagRFP activity measurement

*N. benthamiana* leaves overexpressing SsPrx03-TagRFP, SsPrx03m-TagRFP, and AtPrx36-TagRFP were dissected, weighed (40-60 mg), and frozen in liquid nitrogen. Leaves not transformed or transformed with the pEAQ empty vector were used as negative controls. Tissues were grinded twice for 3 min at 30 Hz using a Retsch MM400 ball mill, with a quick spin between the two cycles. Soluble proteins were extracted in the following extraction buffer (50 ml for 100 mg leaves): 20 mM HEPES, pH 7.0, 1 mM EGTA, 10 mM vitamin C, and PVP PolyclarAT (100 mg⋅g^−1^ fresh weight). The extract was centrifuged twice 10 min at 10,000 g (4°C) to remove insoluble material. The protein content was determined using the Bradford reagent (Serva) with BSA (Sigma-Aldrich) as a standard (Bradford, 1976). CIII Prx activity kinetic was measured at 25°C by following the oxidation of 8 mM guaiacol (Fluka) at 470 nm in the presence of 2 mM H_2_O_2_ (Carlo Erba) in a phosphate buffer (200 mM, pH 6.0). Results were expressed in nanokatals / mg proteins ± SD (n ≥ 4).

## Acknowledgments

The authors are thankful to the Paul Sabatier Toulouse 3 University and to the Centre National de la Recherche Scientifique (CNRS) for granting their work. QL was supported by PhD grants from the China Scholarship Council (CSC). The authors especially thank the Delwiche Lab for the unpublished sequence data available for our analysis and Aurélie Le Ru (UMR5546 CNRS/UPS, Castanet-Tolosan, France) for their valuable help in performing and imaging the transient expression experiments.

**Sup Fig 1 Illustration of cultivated cultures.** Organisms other than the land plants used for the enzymatic assays of the CCP, APx and CIII Prx activities. These organisms are from the Culture Collection of Algae and Protozoa (CCAP, https://www.ccap.ac.uk/). Scale bar = 20 μm.

**Sup Fig. 2. The sequence and structure of SsPrx03.** The positions of the 5 introns are marked with blue down arrows. The Cys residues are marked and numbered as C1, C4 to C8. The Cys residues with the same colour form a disulfide bond (DB): C1 and C4 (red) form DB1; C2 and C3 form DB2, C5 and C8 (blue) form DB3; C6 and C7 (yellow) form DB4. Residues with a green star on top (residue ‘A’ and residue ‘V’) are those mutated in AtPrx36m3 and in. The green helixes under the residue sequence represent the 11 alpha-helix domains and their relative positions in this protein. Key distal and proximal His residues are written as Hd and Hp which are highly conserved.

**Sup Table 1 CI, CII, CIII taxonomic distribution in organism cultivated.** Enumeration of the different peroxidase classes and subclasses in the organisms for which activities were measured (Figure 3). Enumeration was performed only with sequences recorded in PeroxiBase. The absence of an organelle-specific sequence (chloroplast or mitochondria), despite its presence is due to a lack of complete sequencing of certain genomes.

**Sup Table 2 Culture conditions of the different organisms analysed.**

All organisms have been provided by CCAP excepted Moneuplotes crassus and Phaeodactylum Tricornutum which were provided by Professor SWART and Dr. DABOUSSI respectively. PP: Proteose Peptone. MWC: Modified WC Medium. DM: Diatom Medium. PG: Peptone Glucose. PPY: Proteose Peptone Yeast Extract Medium. 3N-BBM+V: Bold Basal Medium with 3-fold Nitrogen and Vitamins.

